# With Childhood Hemispherectomy, One Hemisphere Can Support--But is Suboptimal for--Word and Face Recognition

**DOI:** 10.1101/2020.11.06.371823

**Authors:** Michael C. Granovetter, Sophia Robert, Leah Ettensohn, Marlene Behrmann

## Abstract

The right and left cerebral hemispheres are important for face and word recognition, respectively—a specialization that emerges over human development. The question is whether this bilateral distribution is necessary or whether a single hemisphere, be it left or right, can support both face and word recognition. Here, face and word recognition accuracy in patients with a single hemisphere following childhood hemispherectomy was compared against matched typical controls. In Experiment 1, participants viewed stimuli in central vision. Across both face and word tasks, accuracy of both left and right hemispherectomy patients, while significantly lower than controls’ accuracy, averaged above 80% and did not differ from each other. To compare patients’ single hemisphere more directly to one hemisphere of controls, in Experiment 2, participants viewed stimuli in one visual field to constrain initial processing chiefly to a single (contralateral) hemisphere. Whereas controls had higher word accuracy when words were presented to the right than to the left visual field, there was no field/hemispheric difference for faces. In contrast, left and right hemispherectomy patients, again, showed comparable performance to one another on both face and word recognition, albeit significantly lower than controls. Altogether, the findings indicate that a single developing hemisphere, either left or right, may be sufficiently plastic for comparable representation of faces and words. However, perhaps due to increased competition or “neural crowding,” constraining cortical representations to one hemisphere may collectively hamper face and word recognition, relative to that observed in typical development with two hemispheres.

**Significance Statement:** Adults show right and left cerebral hemispheric biases for face and word recognition, respectively, a division of labor that emerges over development. Here, face and word recognition were assessed in childhood hemispherectomy patients to study the consequences of development with a single hemisphere. While these patients showed above 80% accuracy on face and word recognition tasks, which is surprisingly high relative to the brain volume resected, they nonetheless performed more poorly than typically developing controls. Importantly, patient performance was independent of which hemisphere was removed, suggesting that their single, preserved hemisphere subserves face and word recognition comparably, albeit somewhat inferiorly relative to controls. This demonstrates the remarkable plasticity of the developing brain but, at the same time, highlights plasticity’s constraints.

## Introduction

To the naked eye, anatomical differences between the two cerebral hemispheres of the human brain are largely imperceptible. Decades of data, however, attest to substantial functional differences, for example, with the lateralization of language to the left hemisphere (LH) in the majority of the population (1, 2). A pair of functions with well-established lateralization is face and word recognition, and these hemispheric biases for faces and words in right and left ventral occipitotemporal cortex (VOTC), respectively, appear entrenched by adulthood (3–5). Cortical representations for each stimulus category occupy proximal (and in some cases, overlapping) locations in VOTC, with a weighted asymmetry: greater face selectivity in the right hemisphere (RH) and word selectivity in the left hemisphere (LH) (6–8). Consistently, adults are better at recognizing faces and words when stimuli are presented in a single left or right visual field, respectively, biasing initial processing to one hemisphere over the other (9). In fact, neuropsychological studies in adults suggest that each hemisphere may be *necessary* for the perception of the “preferred” stimulus type: adults with circumscribed focal lesions in left VOTC show significant impairments in word reading (10–12), whereas those with circumscribed focal lesions in left VOTC show significant impairments in face recognition (13–15) (but for deficits in both visual classes, see Behrmann & Plaut (16) & Rice et al. (17)). These sometimes profound behavioral impairments following unilateral lesions lend credence to the claim of relative hemispheric segregation of face and word recognition, at least in the *adult* human brain (18).

Recent investigations of these hemispheric biases for faces and words suggest that they emerge over development via competitive processes for representation in homologous cortex in each hemisphere (19–21). Representation for faces is posited to be represented bilaterally initially in early childhood, although some right-sided bias may be present early on (22). Then, with reading acquisition comes the pressure to optimize proximity of orthographic representations to language regions which are typically left-lateralized. Consequently, word representations become optimized in left VOTC (8, 20, 23) and, by virtue of competition, face representations become optimized in right VOTC. The result is that face and word representations, which become increasingly refined over development, are largely subserved by the RH and LH, respectively (20, 24). Such an account is supported by both behavioral and neuroimaging investigations. For example, in Dundas et al. (9), children and adults discriminated between two faces or two words (in separate blocks). Stimuli were presented briefly in either the participants’ left or right visual fields (with equal probability), thereby restricting initial visual processing to the RH or LH, respectively. A LH over RH performance advantage was evident for words at a younger age than a RH over LH advantage for faces, which was present only in adults. Likewise, using longitudinal functional magnetic resonance imaging (fMRI) in children, Nordt et al. (8) observed increases in word-selectivity in left, but not right, VOTC, with increasing age. They also observed no obvious lateralization for faces in their sample of child participants, even though, by adulthood, fMRI does reveal a RH face bias (25). Meanwhile, Feng et al. (26) reported more word-selectivity in left VOTC in children as a function of reading experience, coupled with a slow, age-related maturation of face-selectivity in right VOTC.

Given 1) the weighted lateralization of these functions, 2) the marked impairment of face or word recognition reported after focal unilateral lesion in adulthood, and 3) the refinement of neural representations for these stimulus categories over development and/or experience (8, 27, 28), the question here concerns the consequences of unilateral hemispheric resection on the lateralization of face and word recognition processes in *children* when the brain is potentially more malleable (29–31). Might a single hemisphere suffice? If so, is this the case irrespective of *which* of the two hemispheres is preserved?

To explore the extent of plasticity, face and word recognition abilities were tested in patients who had undergone childhood hemispherectomy, or the almost complete removal of an entire cerebral hemisphere. One plausible hypothesis is that once literacy begins to be acquired and competition for faces/words has commenced in the developing LH, these representations may continue to emerge with biased lateralization across the two hemispheres, even following surgery. On this account, childhood hemispherectomy patients may show substantial impairments in face or word performance as a function of the removed hemisphere, relative to typically developing controls. That is, left and right hemispherectomy patients would exhibit word and face recognition deficits, respectively, as is the case with adult focal lesion patients. However, given the predisposition for plasticity in childhood (29, 30, 32), an alternative hypothesis is that the competition between faces and words may continue within whichever single hemisphere (LH or RH) is available post-surgery. In this latter case, if face and word representations can emerge within a single hemisphere, competent recognition of both stimulus categories might be observed. Such a finding would attest to substantial plasticity of the human cortex.

The objectives of the current study were thus to determine: 1) whether pediatric patients who develop with only one hemisphere exhibit normative face and word recognition, and 2) whether performance is contingent on which hemisphere is preserved.

## Results

Patients who underwent hemispherectomies as children (here and throughout, the term includes hemispherotomy patients; see Table S1 for patient details) and age- and gender-matched controls viewed pairs of stimuli (pairs of faces or pairs of words, in separate blocks of trials) with sequential presentation of each member of the pair. Participants indicated by pressing one of two keys whether the two stimuli were the same or different (see Materials and Methods and Figure 1 for details). In Experiment 1, both stimuli of the pair were presented foveally (see Fig. 1A). Here, central visual acuity is held constant across groups, but the comparison is between controls who could utilize both hemispheres for visual recognition, compared with patients who could utilize only one hemisphere. In contrast, in Experiment 2, participants viewed the first stimulus of the pair centrally but the second stimulus in one hemifield to restrict initial processing to one hemisphere (see Fig. 1B) (33). This permitted an examination of the approximate competence of a single hemisphere across the two groups by comparing accuracy of a patient’s single hemisphere to a control’s single hemisphere. Trial accuracy (the binary response) was the primary variable of interest, as reaction time (RT) may be confounded by motor impairments in the patients (34, 35).

**Figure 1.**
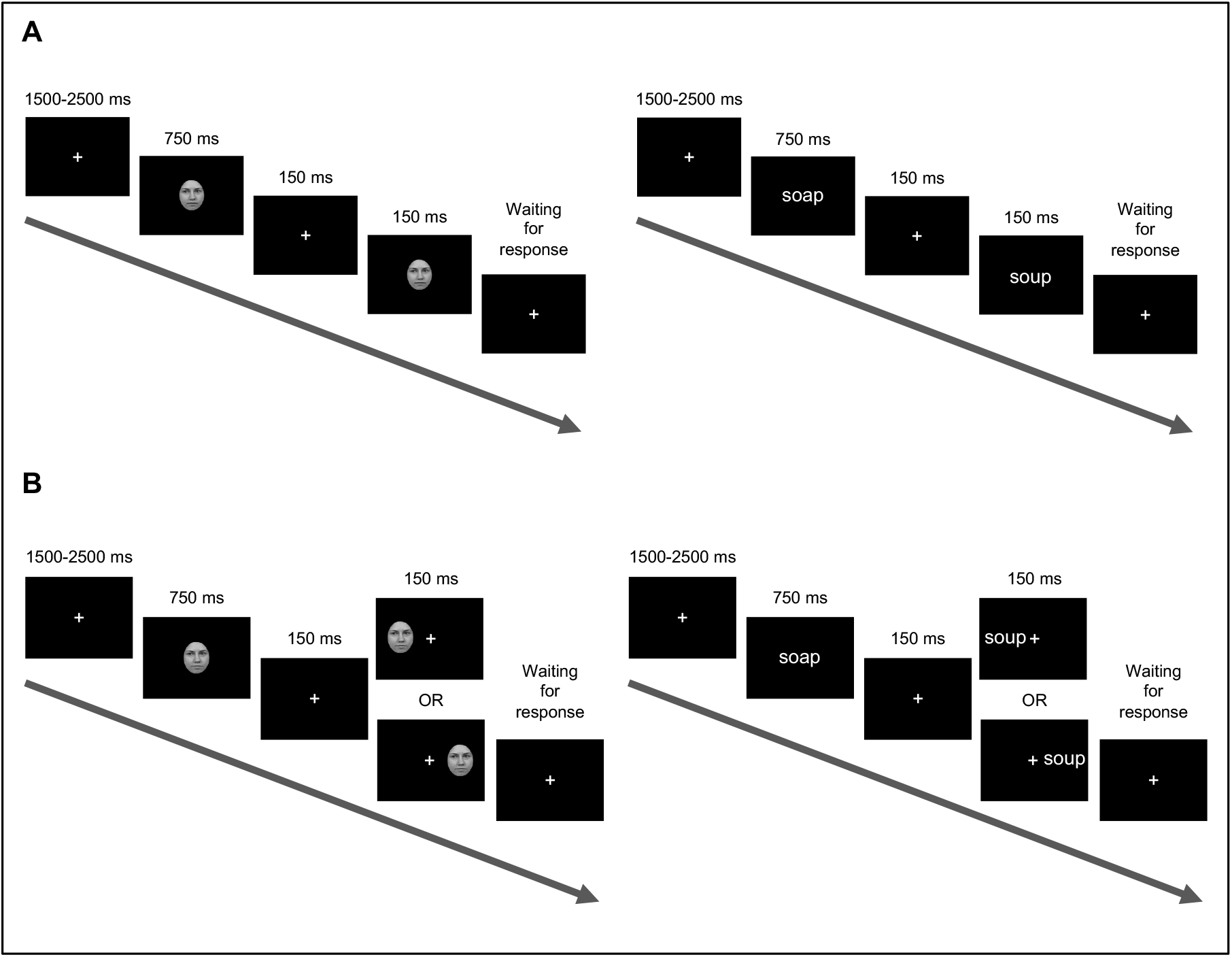
Participants viewed sequential pairs of faces and words (not drawn to scale here) in separate blocks. Participants were instructed to indicate, via one of two response keys, whether stimuli in a pair were the same or different. **A:** In Experiment 1, participants viewed all stimuli at central fixation. **B:** In Experiment 2, participants viewed the second stimulus in one of two visual hemifields to initially restrict processing to controls’ single hemisphere (33), to compare to patients’ preserved single hemisphere.

Furthermore, throughout the text, patients with a preserved LH will be referred to as “LH patients” and patients with a preserved RH as “RH patients.” Two participants had their hemispherectomies completed/revised as adults but were included in the analyses given that their surgeries were initiated in childhood. Differences with versus without these two participants are noted throughout.

### Experiment 1

Data were collected and analyzed from 15 LH patients (median age = 17.9 years, median absolute deviation (MAD) of age = 5.8 yr), 24 RH patients (median age = 15.3, MAD of age = 6.5 yr) and 58 age-matched controls (median age = 17.5 yr, MAD of age = 7.2 yr). A generalized linear mixed effects model (LMEM) was fit to the data (plotted in Fig. 2; see Table S2 for model selection details). Group (controls vs. LH patients vs. RH patients) and stimulus category (faces vs. words), were modeled as predictors of accuracy. Age was also modeled as a covariate, given that hemispheric biases for face and word recognition change over development (20, 36, 37). As a subset of participants were tested online (due to the coronavirus pandemic), whether a testing session was in-person or online was modelled as an additional covariate. Finally, participant was the sole random effect term in each model, with a different intercept per participant.

**Figure 2.**
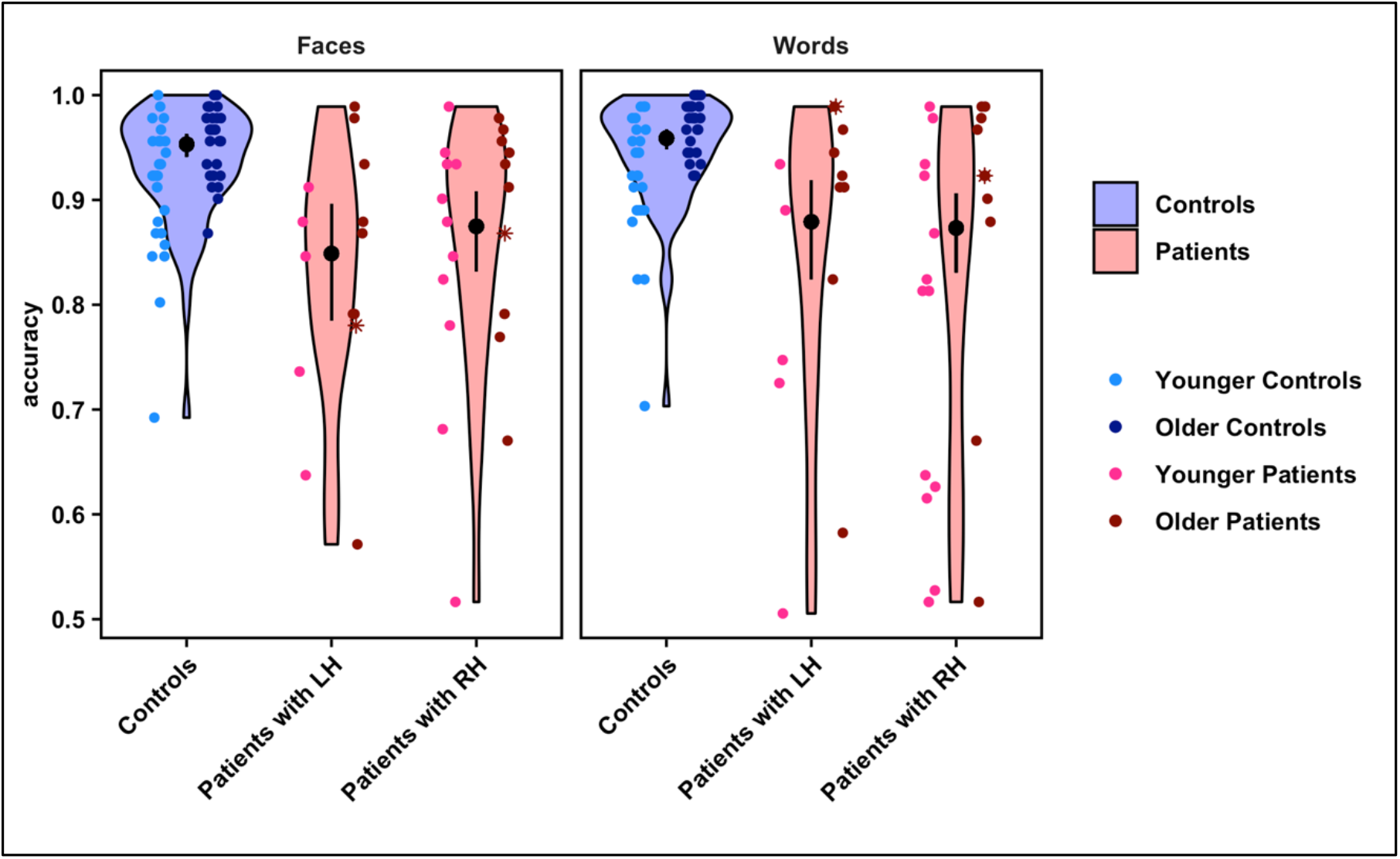
Violin plots show the distribution of overall accuracy values on the face and word recognition tasks for each group and stimulus category. Overlaid point plots show the individual values for each participant. The black points show the estimated probability of correct trials per condition, and the error bars represent the 95% confidence interval of these estimates (back-transformed from a logarithmic scale). To visualize any effect of age, individual participants’ points are separated by the median of the age distribution for all participants in the study (younger versus older than 17.5 yr), shown in different shades of blue (controls) and red (patients). Asterisks indicate the data for two patients in the sample who had their hemispherectomies completed/revised as adults. Points are randomly jittered to minimize overlapping data. Note: axes begin at accuracy = 0.5 because cases in which a participant performed at or below chance on a task were excluded from the analysis.

There was a significant main effect of group on accuracy (*χ*^2_2_^ = 45.97, *p* < 0.001), such that controls performed better than patients on both face and word recognition. Controls had a significantly higher probability of making correct responses (95% confidence interval (CI) = 0.95 - 0.97) than either LH patients (*z* = 5.12, *p* < 0.001; 95% CI = 0.81 - 0.91) or RH patients (*z* = 5.74, *p* < 0.001; 95% CI = 0.83 - 0.91). Furthermore, there was no significant difference between LH and RH patients’ accuracies (*z* = 0.30, *p* = 0.76). This suggests that patients not only showed a deficit on both face and word recognition, but also, importantly, the deficit was independent of the side of resection. At the same time, the average difference in accuracy between patients and controls was no more than 10%, and patients’ accuracy was above 80%. Lastly, there was no significant effect of stimulus category on accuracy (*χ*^2_1_^ = 3.69, *p* = 0.05): performance on one task did not differ from performance on the other, across participants (see Table S3 for model summary and summary statistics).

### Experiment 2

Experiment 1 allowed for a characterization of the behavior of patients with only one hemisphere in comparison to controls with two hemispheres. But an additional question is whether performance of a patient’s single hemisphere is equivalent to the single corresponding--or perhaps, contralateral--hemisphere of a control. Therefore, Experiment 2 used a half-field paradigm in which participants again discriminated between pairs of stimuli, but viewed the second stimulus of each pair in one hemifield to restrict initial processing to the contralateral hemisphere (see Fig. 1B) (33).

In this experiment, 26 hemispherectomy patients participated, all but one of whom also participated in Experiment 1. 11 were LH patients (median age = 18.5 yr, MAD of age = 6.3 yr), and 15 were RH patients (median age = 18.4 yr, MAD of age = 7.6 yr). Patients viewed the second stimulus in each pair in their intact hemifield (the patients are hemianopic) (34). In addition, 15 controls were assigned to view stimuli in their right hemifield (“LH controls”; median age = 18.8 yr, MAD of age = 4.6 yr) and 16 independent controls were assigned to view stimuli in their left hemifield (“RH controls”; median age = 14.6 yr, MAD of age = 5.6 yr). All controls also participated in Experiment 1. A generalized LMEM was fit to these data (plotted in Fig. 3; see Table S4 for model selection details and summary statistics). Group (patients vs. controls), primary hemisphere used (LH vs. RH), stimulus category (faces vs. words), and all interactions between these variables were modeled as predictors of accuracy. As before, age was modeled as a covariate, and participant as a random intercept. (Here, all data were collected online.)

**Figure 3.**
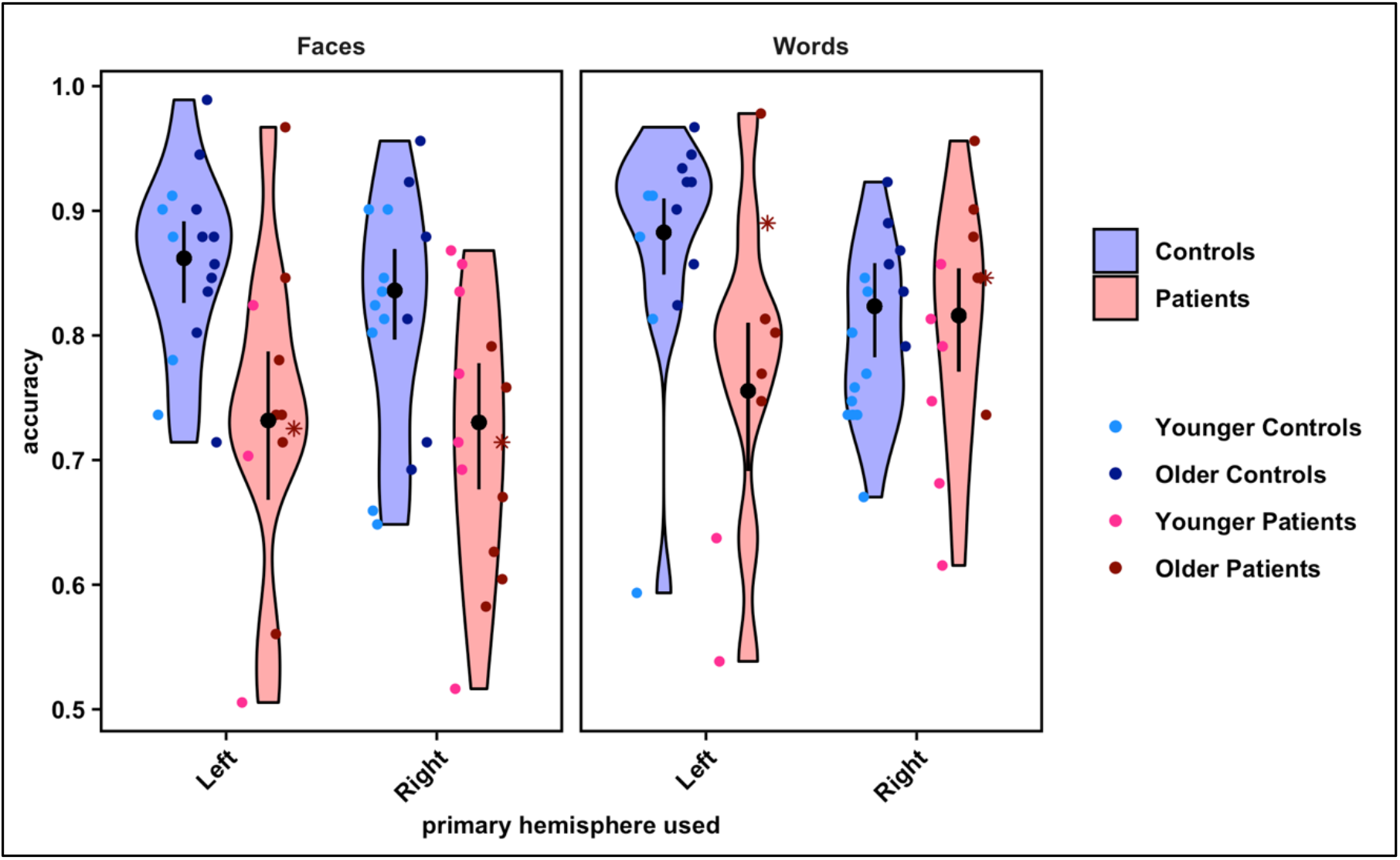
Violin plots show the distribution of overall accuracy values on the face and word recognition tasks for each group and stimulus category, by primary hemisphere used. Overlaid point plots show the individual values for each participant. The black points show the estimated probability of correct trials per condition, and the error bars represent the 95% confidence interval of these estimates (back-transformed from a logarithmic scale). To visualize any effect of age, individual participants’ points are separated by the median of the age distribution for all participants in the study (younger versus older than 17.5 yr), shown in different shades of blue (controls) and red (patients). Asterisks indicate the data for two patients in the sample who had their hemispherectomies completed/revised as adults. Points are randomly jittered to minimize overlapping data. Note: axes begin at accuracy = 0.5 because cases in which a participant performed at or below chance on a task were excluded from the analysis.

Notably, there was a significant three-way interaction of group by hemisphere by stimulus category on accuracy (*χ^2_1_^* = 8.40, *p* < 0.01; see Materials and Methods for model comparison details). Comparing patients to controls, post hoc contrasts demonstrated that LH controls significantly outperformed both patient groups on both tasks. On faces, LH controls’ 95% CI for accuracy was 0.83 - 0.89, while LH patients’ accuracy was 0.67 - 0.79 (*z* = 4.00, *p* < 0.001), and RH patients’ accuracy was 0.68 - 0.78 (*z* = 4.36, *p* < 0.001). On words, LH controls’ 95% CI for accuracy was 0.85 - 0.91, while LH patients’ accuracy was 0.69 - 0.81 (*z* = 4.04, *p* < 0.001), and RH patients’ accuracy was 0.77 - 0.85 (*z* = 2.59, *p* = 0.02). RH controls also significantly outperformed both patient groups, but only on faces, with an accuracy of 0.80 - 0.87 (against LH patients: *z* = 2.99, *p* < 0.01; against RH patients: *z* = 3.40, *p* < 0.01). That is, there was no significant difference in performance on words between RH controls (95% CI = 0.78 - 0.86) and either LH patients (*z* = 1.91, *p* = 0.10) or RH patients (*z* = 0.26, *p* = 0.85). (However, note that after removal of two participants in the sample whose hemispherectomies were completed as adults, RH controls then statistically outperformed LH patients on words, as well; *z* = 2.31, *p* = 0.04.)

Comparing differences between the LH and RH within just the controls, LH controls were significantly more likely to perform better on words than RH controls (*z* = 2.39, *p* = 0.03), but there were no differences between LH and RH controls on faces (*z* = 1.03, *p* = 0.37). These findings are largely consistent with the literature suggesting stronger LH biases for words and weaker RH biases for faces throughout development (9, 20, 21, 37), thereby validating the experimental paradigm. In contrast, comparing just the patients, LH and RH patients showed no differences between each other in performance on *either* task (faces: *z* = 0.04, *p* = 0.97; words: *z* = 1.66, *p* = 0.15). In terms of within-subject comparisons of the effect of task, both LH and RH controls showed no significant *within-subject* differences in performance between the face and word tasks (LH controls: *z* = 1.53, *p* = 0.18; RH controls: *z* = 0.92, *p* = 0.41). LH patients also did not show a within-subject difference between face and word recognition (*z* = 1.05, *p* = 0.37), but RH patients were significantly more likely to discriminate words better than faces (*z* = 4.68, *p* < 0.001). (See Table S5 for model summary and summary statistics.)

Altogether, these results suggest that even when controls are restricted to predominantly using one hemisphere, they largely still outperform patients on face and word recognition, independent of patients’ resection side. One notable exception is that controls restricted to using primarily their RH for word recognition perform comparably to both left and right resection patients. This implies that a patient’s single hemisphere is as disadvantaged for word recognition as a control’s RH, which does not have a preferential bias for word recognition. Furthermore, despite the traditional LH and RH biases for words and faces observed in controls here, side of resection strikingly appears not to affect patients’ task performance. In fact, RH patients have better performance on words than on faces, suggesting that the RH can support typical LH function in the absence of typical LH development.

### Validation Across Experiments

If the results above are reliable and consistent for the hemispherectomized patients, for each individual participant, the extent of impairment should be consistently reproducible across the two experiments. Thus, to validate the reliability of participants’ performance, rank correlations between mean accuracy on Experiment 1 and Experiment 2 were examined, separately for the face and the word recognition tasks (data plotted in Fig. 4). For consistency, correlations were computed (Kendall’s *τ)* across both patient groups (groups were combined given the lack of a significant difference in accuracy on either task between them). Correlations were computed for controls as well, for comparison. *p*-values for the four correlations (two tasks for two groups) were adjusted with the Benjamini & Hochberg correction (38). For the patients, mean accuracy on Experiment 1 was indeed significantly positively correlated with that on Experiment 2 for both the face task (*n* = 23, *τ* = 0.52, *z* = 3.37, *p* < 0.01) and word task (*n* = 20, *τ* = 0.56, *z* = 3.35, *p* < 0.01). For the controls, significant, positive correlations between mean accuracy on Experiments 1 and 2 were also observed, for both the face task (*n* = 30, *τ* = 0.35, *z* = 2.58, *p* = 0.01) and word task (*n* = 29, *τ* = 0.42, *z* = 3.05, *p* < 0.01). In other words, participants’ individual visual recognition performance was reliably reproducible independent of the specifics of the experimental paradigm, attesting to the robustness of the findings.

**Figure 4.**
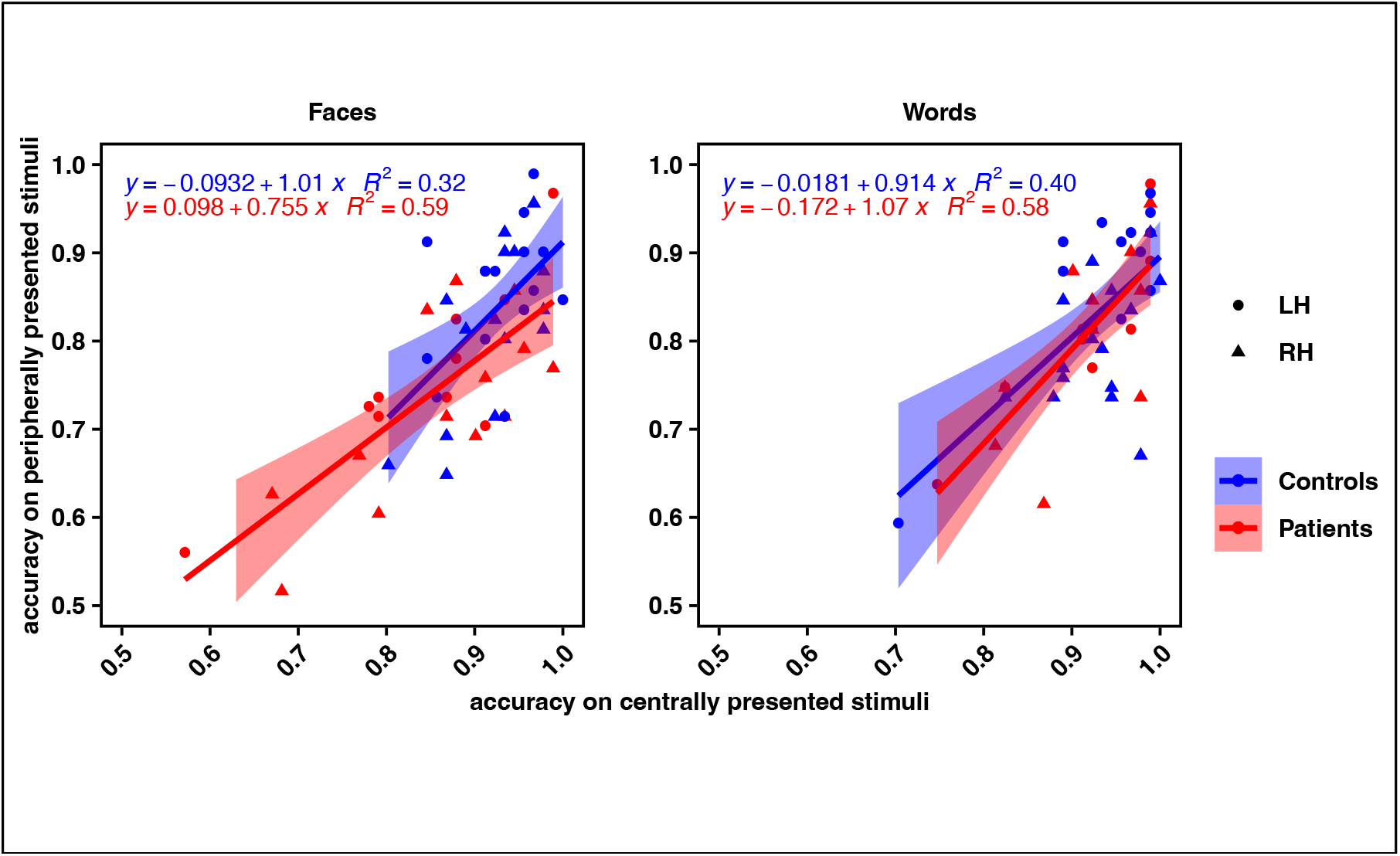
Correlation plots displaying each participant’s accuracy on Experiment 2 (peripheral stimulus presentation; y-axis) versus accuracy on Experiment 1 (central stimulus presentation; x-axis) for faces (left) and words (right). The regression line of best fit for each group is overlaid, with 95% confidence intervals.

### Effects of Clinical Variables

While both LH and RH patients largely performed less accurately on both face and word recognition relative to controls, there is nonetheless heterogeneity in performance across the patients (as in Fig. 2 and 3), as is also evident in Pinabiaux et al. (39). What specific variables might account for this variability? Fig. 5 reveals the performance of individual patients as a function of select variables of interest (collected via parental report). For instance, it is reasonable to consider that the face and word recognition deficits might not be specific to visual recognition per se but, rather, might be attributable to a global cognitive impairment. However, Fig. 5A highlights the heterogeneous cognitive ability of the study’s patient sample and the lack of any apparent relationship with face and word recognition performance. Additionally, ongoing post-surgical seizures may impair plastic processes that would permit one hemisphere to develop both intact face and word recognition. However, the data plotted in Fig. 5B by seizure outcome suggest that the presence of seizures does not obviously account for face and word recognition deficits. A further possibility is that earlier surgery, prior to the establishment of cortical representations for face and word stimuli, would presumably be correlated with better face and word recognition performance. However, since performance increased with age, and younger patients were more likely to have had earlier surgeries, this question could not be formally tested with the current sample, although as evident in Fig. 5C, there does not appear to be an obvious relationship between performance and age at surgery.

**Figure 5.**
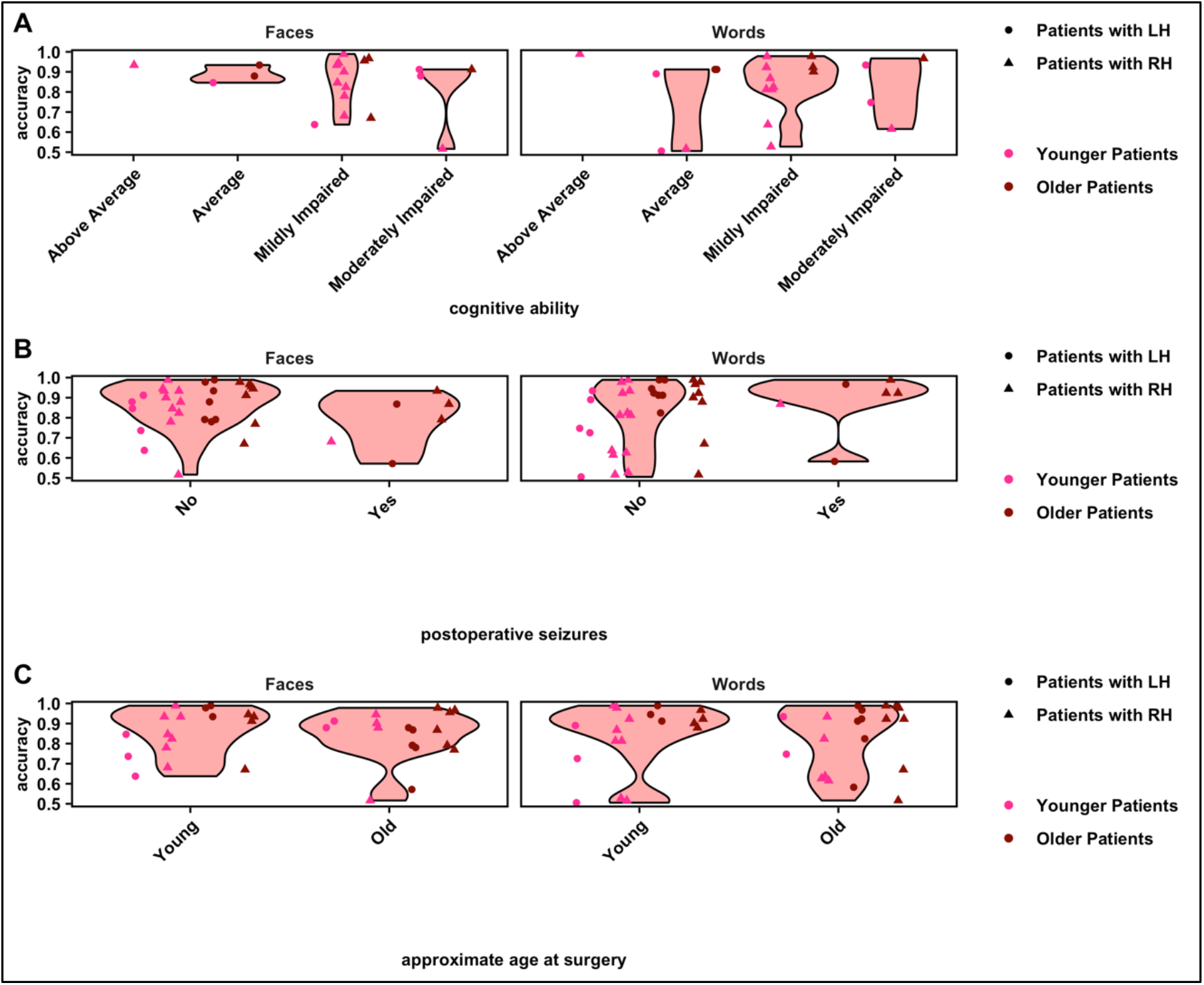
Violin plots show the distribution of overall accuracy values on the face and word recognition tasks for patients, separated by levels of conceivable predictors (per participant and/or guardian report). **A**: Cognitive ability. **B**: Occurrence of any seizures after patients’ most recent surgery. **C**: Approximate age *at surgery* (5 years and younger versus 6 yr and older). Overlaid point plots show the individual values for each participant. To visualize any effect of age *at the time of testing of the study*, individual participants’ points are separated by the median of the age distribution for all participants in the study (younger versus older than 17.5 yr), shown in different shades of red. Points are randomly jittered to minimize overlapping data. Note: axes begin at accuracy = 0.5 because cases in which a participant performed at or below chance on a task were excluded from the analysis.

## Discussion

Canonically, the RH and LH are considered to play important (albeit not exclusive) roles in face and word recognition, respectively (20, 21). However, these hemispheric profiles emerge over development (8, 9, 27), raising the question of whether development with only a single hemisphere suffices for the recognition of faces and words. Furthermore, is this process equivalent if the single hemisphere is the left or the right?

In this study, a relatively large group of hemispherectomy individuals who had previously undergone cortical resection surgery during childhood for pediatric epilepsy treatment was tested on tasks of face and word recognition. The performance of these patients was compared to that of matched controls in two experiments, one in which stimuli were viewed at central fixation (thereby allowing controls to deploy both hemispheres vs. patients’ single hemisphere) and the other in which stimuli were presented in one hemifield (thereby approximately equating groups on a single hemisphere, which receives information from the contralateral field). Three principal findings were revealed across experiments. First, while most patients recognized faces and words with above-chance accuracy (averaging above 80% accuracy under central viewing conditions), their ability to recognize each stimulus category was nonetheless statistically inferior to that of controls’. This performance level is perhaps surprisingly high relative to the brain volume resected (often close to 50%). Second, the patients’ accuracy was not dependent on the hemisphere removed. That is, *the single* LH or RH patients showed comparable performance on face and word recognition. This is true even when participants viewed stimuli in one hemifield, allowing an inferred comparison of the integrity of patients’ single hemisphere to a single hemisphere of controls. It should be noted that the group differences were modulated by the hemisphere (LH, RH) and by the stimulus category (faces, words). Controls using primarily their LH performed better on words than controls primarily using their RH, and there were no differences between LH and RH controls on face accuracy. This result validates the paradigm, as it is consistent with prior results demonstrating the emergence of left-lateralization for words in healthy children, typically prior to right-lateralization for faces (8, 9, 40). In contrast, LH and RH patients performed comparably to each other on both face and word recognition. Both patient groups performed more poorly than controls, with the one exception that they performed comparably to RH controls on words. However, this is consistent with the fact that controls are expected to perform poorer on word recognition when restricted to using the RH (9). Moreover, the results of the two experiments were highly correlated, indicating that the performance metrics for participants remained consistent independent of the specific experimental paradigm. This reliability bolsters the claim that the behavioral profile and upper bound of just a single hemisphere have been rigorously characterized here. Performance levels across patients, while reliably reproducible, were nonetheless heterogenous, but not obviously accounted for by cognitive ability, seizure outcome, or approximate age at surgery.

Altogether, the findings suggest that the preserved hemisphere following childhood hemispherectomy, be it the LH or RH, supports both face and word recognition, albeit inferiorly compared to controls. That is, the typical competition for face and word representations that occurs over normative development in the LH may occur in either the LH or RH of hemispherectomy patients, and constraining the neural competition for representation to a single hemisphere may explain patients’ suboptimal performance relative to healthy participants.

That the pediatric hemispherectomy participants obtain above 80% accuracy on face/word recognition, independent of resection side, is remarkable considering the specific, sometimes profound, deficits that *adult* neuropsychology patients experience after focal VOTC lesions (but see Rice et al. (17)). Numerous case reports and studies have made clear that adults with a left focal (and even, small) lesion are dramatically impaired at word-reading (10–12). Additionally, those with a right focal lesion (even as small as 990 mm^3^) (41) are impaired at face recognition (42, 43). Quantifying these acquired impairments, Behrmann and Plaut reported that adults with a focal LH lesion took roughly 100 times longer (replicated on two separate experiments) to read words of increasing lengths than matched non-neurological controls. In the same study, after a focal RH lesion, adults made roughly ten times more errors than matched controls on two different experiments of face recognition (16). These neuropsychological findings are indeed compatible with experiments in typical adults who demonstrate LH and RH superiorities for word and face recognition, respectively (25, 44, 45).

The lateralized superiorities of the LH for words and the RH for faces appears to emerge over development and are not fully mature in *children* (8, 27, 28). In childhood, it is conjectured that faces are initially represented bilaterally (40). Only over time, as a result of the pressure associated with literacy acquisition, does word representation come to compete with faces for representation in the LH (8, 36, 37). If the representations that typically emerge in the resected hemisphere were instead to arise in the “less-preferred” hemisphere (for example, words in RH after left hemispherectomy), that hemisphere may be optimized for one visual class and, hence, unable to support additional representations. In such a scenario, patients without a LH and patients without a RH would show specific deficits in words and faces, respectively. In contrast, the findings here indicate the recruitment and optimization of the less robust representations in either preserved hemisphere once the “preferred” hemisphere (e.g., LH for words) is no longer functional.

The clear finding is that loss of half of typically available cortex does not result in a proportionate decrement in function, as patients perform at above 80% accuracy. Moreover, the findings here suggest that emergence and competition for face and word representations following literacy acquisition are not limited to the LH but can also occur in the RH. To date, there have been no studies comparing face and word recognition in the same individuals following hemispherectomy, although some studies have investigated face and word recognition independently. For example, one well-known study suggested that the reading ability of hemispherectomy patients appears to depend on age at surgery (and/or seizure onset). In two cases, both with seizure onset after age 10 yr and surgery at age 15 yr, only the individual with the right hemispherectomy showed good (albeit not age-level) reading, similar to the findings reported here. The individual with a left hemispherectomy had a significant and marked reading impairment (46). The latter profile is inconsistent with this current study’s findings but is easily explained by age: the median age at surgery (with earlier onset of seizures) in this current study’s sample was 5.0 yr (MAD = 4.5 yr; median also 5.0 yr for each LH and RH groups; see Figure 5C). For the majority of the children reported here, reading acquisition presumably occurred post-surgically in the preserved RH and was co-localized with language. Additionally, another recent study reported a more frequent decrement in face recognition individuals following right than left resection, albeit using a different approach (standardized neuropsychological testing, only patients with functional hemispherectomy) (39), rendering the comparison of these results with the findings reported in this current study difficult.

The emergence of word representations in the LH is often attributed to the colateralization with the language network, which is generally left-lateralized (19, 23, 47). Hemispherectomy patients with only a RH are almost all verbal to some extent and typically evince post-operative language representation in the RH (48–50). At the same time, functional organization across the two hemispheres is not necessarily discretely segregated. The lateralization of temporal cortex is graded, suggesting that functions that are typically viewed as segregated across the two are nonetheless observable to an extent in each hemisphere (16, 20, 51). Therefore, it follows that the same co-localization of word recognition and language and ensuing competition for words vs. faces may occur in a single preserved RH may be contingent on the age and side of resection of an individual.

Importantly, the patients’ performance was still suboptimal relative to controls. Therefore, while competition between face and word representations may persist following childhood hemispherectomy, the single hemisphere appears inefficient, relative to two hemispheres in typical development, at representing both stimulus categories. Whereas loss of a hemisphere during development does not necessarily interrupt competition between faces and words, it may limit the amount of cortical territory ultimately available for functional specialization. Thus, performance may have an upper bound as a consequence of anatomical constraints: even if the developing brain is sufficiently malleable to allow for emerging face and word representations in one hemisphere, “neural crowding” (32, 52) may hinder the establishment of these representations. This neural competition has in fact been observed in children with more circumscribed resections: children with VOTC resections evince co-existent neural representations for faces and words in the unresected hemisphere, independent of the side of resection (53). Indeed, such competition can be observed longitudinally: a patient with a right VOTC resection showed increasing competition for face representation in the LH over several years (54). Even postoperative changes in functional connectivity in the intact hemisphere of pediatric resection patients may reflect both plasticity and competition for representation of multiple visual categories (55). Future work is needed to probe the direct relationship between behavioral and neural profiles to determine whether the extent of competition for neural representations for faces and words is, in fact, an explanation for patients’ postoperative recognition behaviors. Many other questions--for example, whether the same outcome would result with other visual categories or cognitive functions--can also be addressed.

### Conclusion

Childhood hemispherectomy patients showed above 80% accuracy on tasks of face and word recognition. Despite this surprisingly good performance, it was inferior to that of healthy controls, both when those controls deployed both hemispheres and when they were primarily restricted to using one hemisphere. Importantly, face or word recognition performance was independent of whether the display was of faces or words and independent of the hemisphere resected, and could not be easily explained by cognitive status, extent of seizure freedom, or age at surgery. Together, these results lead to the conclusion that, perhaps via mechanisms of plasticity, either hemisphere appears equally capable of supporting both face and word recognition, although with some cost in performance due to the reduction in available neural substrate. Competition for face and word representations may persist in a sole developing hemisphere. However, given the limited anatomical expanse, neither face nor word recognition can emerge sufficiently to give rise to the competence of visual recognition achievable when the neural labor is divided across the two hemispheres.

## Materials and Methods

### Participants

English-speaking patients with extensive cortical resection (childhood hemispherectomy or hemispherotomy, per participant and/or guardian report; see Kim et al. (34) for distinctions between surgery types) were recruited primarily with the assistance of the Brain Recovery Project: Childhood Epilepsy Surgery Foundation. While right-handers typically evince more reliable LH language lateralization than left-handers (56), native handedness could not be established in patients given their contralesional hemiparesis (34, 35). The sample included a total of 40 patients, 16 with a preserved left hemisphere (“LH patients”; median age = 17.9 yr, MAD of age = 6.1 yr; 12 females, 4 males) and 24 with a preserved right hemisphere (“RH patients”; median age = 15.3, MAD of age = 6.5 yr; 9 females, 15 males; patients’ details are described in full in Table S1). Their performance was compared to that of 58 age-and gender-matched controls (median age = 17.5 yr, MAD of age = 7.2 yr; 30 females, 28 males). These summary statistics reflect the sample used for analysis; sample sizes prior to data quality control (see below for these procedures) are reported in Table S6. Matching each patient group to the control group on age and gender was confirmed with multinomial modelling for Experiment 1; and with logistic modelling for Experiment 2, comparing the patient and control groups for each hemisphere, in separate models. Neither mean-centered age nor gender was significantly predictive of group for either experiment (see Supporting Text for details).

Experimental procedures were approved by the Carnegie Mellon University and University of Pittsburgh Institutional Review Boards. Informed consent was acquired from all adult participants. Child participants gave informed assent, where possible, and guardians provided informed consent. Experiments were conducted either in-person or virtually, with the experimenter(s) and participant (and parent or guardian too, where necessary) communicating via Zoom.

### Stimuli

Stimuli were adopted from Dundas et al. (9) who studied a sample of children and adults of ages comparable to those here.

#### Faces

Grayscale, neutral-expressive, forward-facing 48 unique face stimuli from the Face Place Database (57), edited to remove hair, were presented against a black background.

#### Words

Pairs of 58 four-letter gray words, chosen to be age-appropriate for young readers, were presented in Arial font, on a black background. Words in each pair differed by one letter (for example, “tack” vs. “tank”). The second letter differed in 14 pairs, and the third letter differed in 15.

### Procedures

Per trial, participants viewed one stimulus (either a face or a word, with the categories presented in separate blocks) centrally for 750 ms (long enough for encoding by children). After a 150 ms interval with just a fixation cross, participants saw a second stimulus for 150 ms (too brief to plan and execute a saccade). A randomly jittered 1500 to 2500 ms interval separated trials. Participants reported with a key press, whether the two stimuli were the same or different, for 96 trials (half same, half different). Stimulus pairs were randomized within-block (see Supporting Text for details). In each experiment, participants acclimated to the tasks by completing 12 practice trials prior to the first block of each stimulus category. No direct feedback was provided during the experiment, but general encouragement was provided throughout.

Participants were instructed to maintain central fixation. In Experiment 1, all stimuli were presented centrally (see Fig. 1A). In Experiment 2, the second stimulus in a matching pair was presented in only one hemifield at an eccentricity of approximately 5° from fixation, such that the information would initially be primarily processed by the hemisphere contralateral to the hemifield of presentation (33) (see Fig. 2A). In this Experiment 2, patients, who are hemianopic (34), viewed peripheral stimuli only in their intact hemifield. To match the patient paradigm, two groups of controls of approximately equal size were recruited to view all peripheral stimuli in only either their left hemifield (“RH controls”) or right hemifield (“LH controls”). To encourage the maintenance of central fixation, in addition to the first stimulus in each pair being presented centrally and the second, peripheral stimulus with brief duration, 12 “catch” trials were included per experiment block where the second stimulus was presented centrally instead of peripherally. (See Supporting Text for details on randomization of catch trials into the trial order.)

For in-person sessions, participants viewed stimuli with the Psychophysics Toolbox (58) in MATLAB (MathWorks), and for virtual sessions, participants viewed stimuli in PsychoPy (59) via the Pavlovia interface. In-person participants only participated in Experiment 1, and the majority of them completed the word block before the face block. Most online participants participated in Experiment 2, followed by Experiment 1, with ordering of face and word blocks within each experiment approximately counterbalanced across participants. Only a subset of participants participated in Experiment 2, and some participants did not complete both experiments and/or blocks: sample size details are reported in Table S6.

Some participants participated both in-person and online (*n* = 19). Comparisons of their in-person and online data revealed no significant *within*-subjects effect of session type on accuracy (*χ*^2_1_^ = 0.04, *p* = 0.83) supporting the reliability of individual participants’ data across the presentation formats. For participants who participated both in-person and online, only the online data were ultimately used for analysis. Still, with unique participants participating in-person or online, there was a significant *between*-subjects effect of session type on accuracy (*χ*^2_1_^ = 9.97, p < 0.01; see Supporting Text for details on these comparisons). Thus, session type was modeled as a covariate for analyses including both in-person and online data.

### Data Quality Control

All trials required a response, with no time limit. Thus, trials with RTs longer than the 95th percentile of a participant’s RT distribution on a given experiment block (central or peripheral presentation of faces or words) were discarded. Moreover, if participants were indeed fixating centrally, their performance on trials with peripheral stimuli should not exceed that on catch trials (with only central stimuli). Therefore, data on an experiment block were discarded if the participant’s accuracy was significantly lower or RT was significantly longer on catch than on experimental trials (using permutation testing on models predicting accuracy or RT from trial type; see Supporting Text for details). Data were also discarded if average accuracy across catch trials was at or below 50%. Furthermore, an experiment block was discarded from the analysis if the participant’s accuracy for that block was at or below chance. Below chance accuracy was observed in only nine blocks (across all Experiment and stimulus type combinations) for only six patients. All such cases were patients and represented a minority of the total recruited patient sample. Finally, if mean RT on a block was greater than three standard deviations from the mean of the distribution of mean RTs for the given group (patient vs. control; and hemisphere used, if applicable), data on that block were discarded.

### Data Analysis

Statistical analyses (and data quality control) were conducted in R version 3.6.3 (60) (for complete list of packages, see Table S7). LMEMs were fit as described by Brown (61) and Fox & Weisberg (62). For Experiment 1, a LMEM predicting accuracy was modelled with the two-way interaction--as well as individual main effects--of group (controls vs. LH patients vs. RH patients) and stimulus category (faces vs. words) as fixed effects of interest; mean-centered age and session type (in-person vs. virtual) as fixed covariates; and participant as a random intercept. The model with the interaction term was compared to a null model with no interaction using a likelihood ratio test (LRT). The Akaike information criterion (AIC) of the model without the interaction was lower and afforded a significantly better fit to the data than the model with it. Thus, the interaction term was dropped from the ultimate model.

For Experiment 2, a LMEM predicting accuracy was also modelled, but initially with fixed effects of the three-way interaction--as well as individual main effects--of group (controls vs. patients), hemisphere used (LH vs. RH), and stimulus category (faces vs. words) as fixed effects; mean-centered age as a fixed covariate, and participant as a random intercept. (Note: session type was not modeled as a covariate here because Experiment 2 was only conducted with the online format.) The full model was compared to the model with all three two-way interactions with an LRT. The AIC of the model with the three-way interaction term was lower and afforded a significantly better fit to the data than the model without it, and, therefore, the former was evaluated.

Fixed effects were estimated to optimize the maximum log-likelihood criterion (63) and were fit using the Bound Optimization by Quadratic Approximation (BOBYQA) algorithm. References to age in the Results refer to the mean-centered age (that is, age subtracted from the mean of all participants in an analysis) (64). *p*-values were computed for each model term with Type II Wald chi-square tests (62). Additionally, planned post hoc contrasts were performed by comparing estimated log odds ratios to 1 (for analyses on accuracy) with *z*-tests (65). For each set of planned contrasts, *p*-values were adjusted with the Benjamini & Hochberg correction (38). Estimates and confidence intervals were computed assuming degrees of freedom of infinity. The a threshold for significance was set at 0.05.

Experiment and analysis code, raw data, and stimuli will be available to reviewers upon request during the peer-review process and, upon publication, will be made publicly available on KiltHub (CMU; reserved digital object identifier: 10.1184/R1/12743276).

## Acknowledgments

This research was supported by Award Numbers R01EY027018 from the National Eye Institute (NEI) to MB and T32GM081760 from the National Institute of General Medical Sciences (NIGMS) to MCG. MCG was also supported by a fellowship from the American Epilepsy Society (AES), SR was supported by a fellowship from the National Science Foundation (NSF), and LE was supported by a grant from the Carnegie Mellon University (CMU) undergraduate research office. The content is solely the responsibility of the authors and does not necessarily represent the official views of the NEI, NIGMS, National Institutes of Health, AES, NSF, or CMU. The authors are grateful to the Brain Recovery Project: Childhood Epilepsy Surgery Foundation, especially its founder and Chief Executive Officer Monika Jones, JD, for facilitating patient data collection, and to Tekla Hilton and Avi Munro for facilitating control data collection at Community Day School. The authors are also grateful to Michael J. Tarr, Department of Psychology, Carnegie Mellon University, http://www.tarrlab.org/ for the use of the stimuli from the Face Place database (funding provided by NSF award 0339122). The authors also acknowledge Dr. Marge Maallo for her shared experiment code and input on the study design; Dr. Maallo and Raina Vin for their assistance with data collection; Drs. Carl Olson, David Plaut, and Tarr for their feedback on the study design and data interpretation; Drs. Alison Butler, Joel Greenhouse, and Alexander Layden for their input on statistical analyses; Nicholas Blauch and Drs. Vladislav Ayzenberg, Tina Liu, Maallo, Plaut, and Tarr for their feedback on drafts of this manuscript; and the CMU Visual Cognition group for their helpful discussions. Finally, the authors thank the participants and their families for their cooperation.

