## Supporting Information for "With Childhood Hemispherectomy, One Hemisphere Can Support--But is Suboptimal for--Word and Face Recognition"

**This PDF file includes:**

Supporting text
Tables S1 to S9
SI References

### Supporting Information Text

#### Confirmation that groups were matched on age and gender (after data quality control).

Experiment 1

With data used from Experiment 1 (central stimulus presentation), multinomial models were fit with group (controls vs. LH patients vs. RH patients) as the dependent variable. Mean-centered age was the predictor in one model, gender in another. Age was not significantly predictive of whether a participant was a control vs. either a LH patient ($e^{\beta}$ = 1.03, 95% CI = 0.96 - 1.11, $z_4$ = 0.94, $p$ = 0.35) or a RH patient ($e^{\beta}$ = 1.01, 95% CI = 0.95 - 1.08, $z_4$ = 0.26, $p$ = 0.79). Gender also was not significantly predictive of whether a participant was a control vs. either a LH patient ($e^{\beta}$ = 0.39, 95% CI = 0.11 - 1.37, $z_4$ = 1.47, $p$ = 0.14) or a RH patient ($e^{\beta}$ = 1.79, 95% CI = 0.67 - 4.73, $z_4$ = 1.17, $p$ = 0.24).

Experiment 2

With data used from Experiment 2 (peripheral stimulus presentation where both patients and controls were chiefly using one hemisphere), ages and genders of LH controls were compared to LH patients, and ages and genders of RH controls were compared to RH patients. For each comparison, logistic models were fit with group as the binomial outcome variable, with mean-centered age as the predictor in one model, and gender in the other. Age was not significantly predictive of whether a participant was a patient or control for either LH participants ($e^{\beta}$ = 1.02, 95% CI = 0.92 - 1.13, $z_2$ = 0.39, $p$ = 0.70) or RH participants ($e^{\beta}$ = 1.09, 95% CI = 0.94 - 1.28, $z_2$ = 1.08, $p$ = 0.28). Gender also was not significantly predictive of whether a participant was a patient or control for either LH participants ($e^{\beta}$ = 0.75, 95% CI = 0.12 - 4.06, $z_2$ = 0.33, $p$ = 0.74) or RH participants ($e^{\beta}$ = 1.20, 95% CI = 0.27 - 5.44, $z_2$ = 0.24, $p$ = 0.81).

#### Randomization of stimulus presentations.

Faces

For in-person sessions (96 trials of centrally presented face pairs only), all 48 face stimuli were shown as the first stimulus in a pair (the "target" stimulus) twice each. For trials in which the stimuli within a pair differed, the second stimulus in the pair (the "probe" stimulus) was a randomly designated face from any of the 47 other stimuli. For online sessions, for trials in which the stimuli within a pair differed, each of the 48 target stimuli were paired with a pseudo-randomly selected probe: the randomized list of target stimuli was divided into four lists of 12, and the probe was randomly selected from one of the 12-item lists that did not contain the target stimulus. In this way, on trials in which the stimuli within a pair differed, each of the 48 stimuli were shown as probe stimulus once each, with no matching stimuli within a pair. Furthermore, online participants completed a block with probe stimuli displayed peripherally (using the same randomization scheme described above), but with the addition of 12 catch trials. For these trials, 18 stimuli were randomly selected: 6 for trials in which stimuli were the same, and 12 were paired together for trials in which stimuli differed.

Words

For in-person sessions (96 trials of centrally presented word pairs only), each of the 58 word stimuli were shown as the target stimulus once each in the first 58 trials; in the remaining trials, 38 of the 58 stimuli were randomly selected as the target stimulus. For online sessions, there were two types of stimulus pairs: word pairs either differed in the second letter (14 pairs of 28 stimuli) or in the third letter (15 pairs of 30 stimuli). Twenty-four unique words were randomly selected from each type of word pair once for trials in which stimuli were the same (48 trials), and 24 pairs of matched words were randomly selected from each type of word pair for trials in which stimuli differed (48 trials). The 96 word pairs were then shuffled such that the four trial types were dispersed throughout the run. For those participants viewing stimuli peripherally, 4 unique catch trials composed of word pairs not included in the regular trials were shuffled into each of three mini-blocks of 32 trials. The 12 total catch trials were composed of equal numbers of pairs of stimuli that were the same or different, and equal numbers of pairs of differing stimuli in which the second or third letter differed.

**Comparison of in-person and online accuracy data.** Nineteen participants participated in Experiment 1 both in-person and online. On these data, a generalized LMEM was fit with trial accuracy as the binomial dependent variable, session type as the fixed effect of interest, and mean-centered age as a fixed covariate (of note, in-person and online sessions occurred approximately two yr apart, therefore especially necessitating this covariate). Participant was modeled as a random intercept. A Type II Wald chi-square test demonstrated no significant effect of session type on trial accuracy ($e^{\beta}$ = 0.98, 95% CI = 0.80 - 1.19, $\chi^2_1$ = 0.04, $p$ = 0.83; Table S8).

In addition, on the data used for analysis, in which in-person data were discarded for participants who participated in Experiment 1 both in-person and online, a generalized LMEM was fit again with trial accuracy as the binomial dependent variable, session type as the fixed effect of interest, and mean-centered age as a fixed covariate (although unlike above, here session type is a *between*-, not within-subjects effect). A Type II Wald chi-square test indicated that participants were significantly less likely to provide correct responses online than in-person ($e^{\beta}$ = 0.51, 95% CI = 0.33 - 0.77, $\chi^2_1$ = 9.97, $p$ < 0.01; Table S9), justifying the inclusion of session type as a between-subjects covariate in the analyses reported in the Results.

**Discarding data for trials on which participants may not have been fixating centrally.** In Experiment 2, if a participant was centrally fixating, their accuracy on trials with peripherally presented stimuli ("test trials") should not exceed that on trials with centrally presented stimuli ("catch trials"). To assess whether this was the case, accuracy and RT were each compared on catch vs. test trials. Given the low sample size for catch trials ($n$ = 12), permutation testing was employed. For each participant on each experiment block (faces or words), the trial type label (catch vs. test) was shuffled 1000 times, and, on each permutation, a logistic model was fit with trial accuracy as the dependent variable and trial type as the predictor. The estimated probability that correct responses would be more likely on test vs. catch trials due to chance ($p$) was computed as the sum of instances in which the $\beta$ coefficient of the trial type predictor from a model on the permuted data was greater than that from the same model on the true data. The trial type label was then again shuffled 1000 times on each experiment block, but here, on each permutation, a general linear model was fit with RT as the dependent variable (and trial type again as the predictor). Here, the estimated probability that *faster* RTs would be more likely on test vs. catch trials due to chance was computed as the sum of instances in which the $\beta$ coefficient of the trial type predictor from a model on the permuted data was *less* than that from the same model on the true data. $p$-values across participants were adjusted with the Benjamini & Hochberg correction (1), separately for each block type and dependent measure. Individual participants' individual experiment blocks with an adjusted $p$-value less than 0.05 for the catch vs. test trial comparisons on either accuracy or RT were then discarded.

**Table S1.** Patient information.

| Age (years) | Gender | ~Age at First Surgery (yr)* | Cognitive Ability** | Postoperative Seizure(s) |
|---|---|---|---|---|
| **Left Hemisphere Surgery Cases** | | | | |
| 8.0 | Female | 1 | Mildly Impaired | No |
| 8.1 | Female | 1 | Mildly Impaired | No |
| 8.6 | Male | 1 | Average | No |
| 9.8 | Female | 7 | - | No |
| 10.7 | Female | 5 | Mildly Impaired | Yes |
| 10.7 | Female | 6 | Mildly Impaired | No |
| 10.9 | Male | 1 | Mildly Impaired | No |
| 11.7 | Male | 4 | Mildly Impaired | No |
| 13.1 | Female | 10 | - | No |
| 13.3 | Male | 4 | Mildly Impaired | No |
| 14.1 | Female | 10 | Moderately Impaired | No |
| 15.1 | Male | 5 | Above Average | No |
| 15.4 | Female | 13 | Mildly Impaired | No |
| 18.1 | Male | 6 | - | No |
| 18.4 | Male | 14 | Mildly Impaired | No |
| 18.9 | Male | 4 | Mildly Impaired | No |
| 19.0 | Male | 8 | - | No |
| 19.6 | Male | 7 | Mildly Impaired | No |
| 19.6 | Female | 4 | Moderately Impaired | No |
| 20.6 | Male | 8 | - | Yes |
| 23.7 | Male | 6 | - | No |
| 30.9 | Male | 1* | - | Yes |
| 31.4 | Male | 4 | - | No |
| 38.8 | Male | 1 | - | Yes |
| **Right Hemisphere Surgery Cases** | | | | |
| 6.5 | Female | 3 | Mildly Impaired | No |
| 6.7 | Female | 3 | Average | No |
| 9.1 | Female | 2 | - | No |
| 13.5 | Female | 11 | Mildly Impaired | No |
| 14.0 | Female | 6 | Moderately Impaired | No |
| 14.3 | Female | 13 | Moderately Impaired | No |
| 15.5 | Female | 1 | Average | No |
| 17.9 | Male | 2 | Average | No |
| 17.9 | Female | 9 | Average | No |
| 18.5 | Male | 6 | - | Yes |
| 20.6 | Female | 13 | - | No |
| 20.7 | Female | 12 | - | No |
| 23.6 | Male | 0 | - | No |
| 25.0 | Female | 4 | - | No |
| 31.5 | Female | 8 | - | Yes |
| 37.0 | Male | 11* | - | No |

* Surgery ages are per participant and/or their guardian report. Some patients had multiple surgeries to complete the hemispherectomy, and two of these patients had their surgeries completed as adults. Excluding these participants from the analyses does not affect the primary interpretation of the results. Minor discrepancies in statistical significance with or without these two participants are discussed in the Results.

** Per guardian report. Guardians' cognitive ability assessment is only reported for participants who were younger than 18 yr at the initial time of study participation.

- No information available.

**Table S2.** Model selection for predicting accuracy on Experiment 1 data.

| **Models** | | |
|---|---|---|
| | *Model* | *AIC* |
| *1* | accuracy ~ group * stimulus category + group + stimulus category + age + session type + (1 \| participant) | $9.55*10^3$ |
| *2* | accuracy ~ group + stimulus category + age + session type + (1 \| participant) | $9.55*10^3$ |
| **Likelihood Ratio Test** | | |
| | $\chi^2_2$ | *p* |
| *1 vs. 2* | 3.22 | 0.20 |

**Table S3.** Model summary for Experiment 1 accuracy data.

| **accuracy ~ group + stimulus category + age + session type + (1 \| participant)** | | | |
|---|---|---|---|
| **Type II Wald $\chi^2$ tests** | | | |
| | $\chi^2$ | df | *p* |
| *group* | 45.97 | 2 | < 0.001 |
| *stimulus category* | 3.69 | 1 | 0.05 |
| *age* | 29.35 | 1 | < 0.001 |
| *session type* | 5.61 | 1 | 0.02 |

| **Estimated probabilities of correct accuracy** | | |
|---|---|---|
| *group* | *estimated probability* | *95% CI* |
| Controls | 0.96 | 0.95 - 0.97 |
| LH Patients | 0.86 | 0.81 - 0.91 |
| RH Patients | 0.87 | 0.83 - 0.91 |
| **Post hoc contrasts on group** | | |
| | *z* | *p* |
| *Controls / LH Patients* | 5.12 | < 0.001 |
| *Controls / RH Patients* | 5.74 | < 0.001 |
| *LH Patients / RH Patients* | -0.30 | 0.76 |

**Table S4.** Model selection for predicting accuracy on Experiment 2 data.

| **Models** | | |
|---|---|---|
| | *Model* | *AIC* |
| *1* | acc ~ group * hemisphere * stimulus category +<br>group * hemisphere + group * stimulus category + hemisphere * stimulus category +<br>group + hemisphere + stimulus category +<br>age +<br>(1 \| participant) | $9.19*10^3$ |
| *2* | acc ~ group * hemisphere + group * stimulus category + hemisphere * stimulus category +<br>group + hemisphere + stimulus category +<br>age +<br>(1 \| participant) | $9.19*10^3$ |

| **Likelihood Ratio Test** | | | |
|---|---|---|---|
| | $\chi^2$ | *df* | *p* |
| *1 vs. 2* | 8.37 | 1 | < 0.01 |

acc = accuracy

**Table S5.** Model summary for Experiment 2 accuracy data.

| **accuracy ~ group * hemisphere * stimulus category +<br>group * hemisphere + group * stimulus category + hemisphere * stimulus category +<br>group + hemisphere + stimulus category +<br>age + session type +<br>(1 \| participant)** | | |
|---|---|---|
| **Type II Wald $\chi^2$ tests** | | |
| | $\chi^2_1$ | *p* |
| *group * hemisphere * stimulus category* | 8.40 | < 0.01 |
| *group * hemisphere* | 3.27 | 0.07 |
| *group * stimulus category* | 8.22 | < 0.01 |
| *hemisphere * stimulus category* | 0.13 | 0.72 |
| *group* | 19.79 | < 0.001 |
| *hemisphere* | 0.73 | 0.39 |
| *stimulus category* | 9.54 | < 0.01 |
| *age* | 13.15 | < 0.001 |

| **Estimated probabilities of correct accuracy** | | | | |
|---|---|---|---|---|
| *group* | *hemisphere* | *stimulus category* | *estimated probability* | *95% CI* |
| Controls | LH | Faces | 0.86 | 0.83 - 0.89 |
| | | Words | 0.88 | 0.85 - 0.91 |
| | RH | Faces | 0.84 | 0.80 - 0.87 |
| | | Words | 0.82 | 0.78 - 0.86 |
| Patients | LH | Faces | 0.73 | 0.67 - 0.79 |
| | | Words | 0.76 | 0.69 - 0.81 |
| | RH | Faces | 0.73 | 0.68 - 0.78 |
| | | Words | 0.82 | 0.77 - 0.85 |

| **Post hoc contrasts on group * hemisphere * stimulus category** | | |
|---|---|---|
| | *z* | *p* |
| *Controls LH Faces / Patients LH Faces* | 4.00 | < 0.001 |
| *Controls LH Faces / Patients RH Faces* | 4.36 | < 0.001 |
| *Patients LH Faces / Controls RH Faces* | -2.99 | < 0.01 |
| *Controls RH Faces / Patients RH Faces* | 3.40 | < 0.01 |
| *Controls LH Words / Patients LH Words* | 4.04 | < 0.001 |
| *Controls LH Words / Patients RH Words* | 2.59 | 0.02 |
| *Patients LH Words / Controls RH Words* | -1.91 | 0.10 |
| *Controls RH Words / Patients RH Words* | 0.26 | 0.85 |
| *Controls LH Faces / Controls RH Faces* | 1.03 | 0.37 |
| *Controls LH Words / Controls RH Words* | 2.39 | 0.03 |
| *Patients LH Faces / Patients RH Faces* | 0.04 | 0.97 |
| *Patients LH Words / Patients RH Words* | -1.66 | 0.15 |
| *Controls LH Faces / Controls LH Words* | -1.53 | 0.18 |
| *Controls RH Faces / Controls RH Words* | 0.92 | 0.41 |
| *Patients LH Faces / Patients LH Words* | -1.05 | 0.37 |
| *Patients RH Faces / Patients RH Words* | -4.68 | < 0.001 |

**Table S6.** Sample sizes per session type, experiment, group, and stimulus category block type, pre- and post-data data quality control (see Materials and Methods for descriptions of data quality control steps).

| | Pre-Quality Control Sample Sizes | | | | Post-Quality Control Sample Sizes | | | |
|---|---|---|---|---|---|---|---|---|
| | In-Person Only | Online Only | Both | Total | In-Person Only | Online Only | Both | Total |
| **Experiment 1 (Central Stimulus Presentation)** | | | | | | | | |
| *Faces* | | | | | | | | |
| Controls | 23 | 28 | 3 | 54 | 23 | 28 | 3 | 54 |
| Patients with LH | 3 | 8 | 4 | 15 | 3 | 7 | 4 | 14 |
| Patients with RH | 6 | 11 | 6 | 23 | 6 | 9 | 6 | 21 |
| *Words* | | | | | | | | |
| Controls | 26 | 27 | 3 | 56 | 25 | 27 | 3 | 55 |
| Patients with LH | 3 | 8 | 5 | 16 | 2 | 7 | 5 | 14 |
| Patients with RH | 7 | 7 | 10 | 24 | 7 | 7 | 10 | 24 |
| *Totals* | | | | | | | | |
| Controls | 27 | 27 | 4 | 58 | 27 | 27 | 4 | 58 |
| Patients with LH | 3 | 8 | 5 | 16 | 3 | 7 | 5 | 15 |
| Patients with RH | 7 | 7 | 10 | 24 | 7 | 7 | 10 | 24 |
| **Experiment 2 (Peripheral Stimulus Presentation)** | | | | | | | | |
| *Faces* | | | | | | | | |
| LH Controls | - | 15 | - | 15 | - | 15 | - | 15 |
| RH Controls | - | 16 | - | 16 | - | 15 | - | 15 |
| Patients with LH | - | 13 | - | 13 | - | 11 | - | 11 |
| Patients with RH | - | 17 | - | 17 | - | 14 | - | 14 |
| *Words* | | | | | | | | |
| LH Controls | - | 14 | - | 14 | - | 13 | - | 13 |
| RH Controls | - | 16 | - | 16 | - | 16 | - | 16 |
| Patients with LH | - | 12 | - | 12 | - | 9 | - | 9 |
| Patients with RH | - | 17 | - | 17 | - | 12 | - | 12 |
| *Totals* | | | | | | | | |
| LH Controls | - | 15 | - | 15 | - | 15 | - | 15 |
| RH Controls | - | 16 | - | 16 | - | 16 | - | 16 |
| Patients with LH | - | 13 | - | 13 | - | 11 | - | 11 |
| Patients with RH | - | 17 | - | 17 | - | 15 | - | 15 |

LH = left hemisphere
RH = right hemisphere
LH and RH controls = controls viewing stimuli in their right and left visual fields, respectively, thereby restricting initial processing to the LH and RH, respectively (2).
- Not applicable.

**Table S7.** R packages used for data cleaning and analysis.

| Package | Version | Use |
|---|---|---|
| broom (3) | 0.7.9 | Summarizing statistics |
| car (4) | 3.0-11 | Summarizing statistics |
| emmeans (5) | 1.6.2-1 | Computing estimates and performing post hoc comparisons |
| ggnewscale (6) | 0.4.5 | Plotting |
| ggpmisc (7) | 0.4.5 | Plotting |
| ggpubr (8) | 0.4.0 | Plotting |
| lme4 (9) | 1.1-27.1 | Fitting linear mixed models |
| nnet (10) | 7.3-12 | Fitting multinomial models |
| plyr (11) | 1.8.6 | Manipulating data |
| pracma (12) | 2.3.3 | Manipulating data |
| psych (13) | 2.1.9 | Computing descriptive statistics |
| stringr (14) | 1.4.0 | Manipulating data |
| tidyverse (15) | 1.3.1 | Manipulating data |

**Table S8.** Model summary of the generalized linear mixed model: accuracy ~ session type + age + (1 | participant)

| **Type II Wald $\chi^2$ tests** | | |
|---|---|---|
| | $\chi^2_1$ | *p* |
| *session type* | 0.04 | 0.83 |
| *age* | 5.48 | 0.02 |
| **Exponentiated Fixed Effects and 95% Confidence Bounds** | | |
| | *Estimate* | *95% CI* |
| *session type* | 0.98 | 0.80 - 1.19 |
| *age* | 1.06 | 1.01 - 1.12 |

Note: this model was fit only to data in which participants participated in Experiment 1 both in-person and online.

**Table S9.** Model summary of the generalized linear mixed model: accuracy ~ session type + age + (1 | participant)

| **Type II Wald $\chi^2$ tests** | | |
|---|---|---|
| | $\chi^2_1$ | *p* |
| *session type* | 9.97 | < 0.01 |
| *age* | 17.06 | < 0.001 |
| **Exponentiated Fixed Effects and 95% Confidence Bounds** | | |
| | *Estimate* | *95% CI* |
| *session type* | 0.51 | 0.33 - 0.77 |
| *age* | 1.06 | 1.03 - 1.09 |

Note: this model was fit only to data used for the analyses in Results; in-person data were discarded for all participants who participated both in-person and online.
